# Synthesis of geological and comparative phylogeographic data point to climate, not mountain uplift, as driver of divergence across the Eastern Andean Cordillera

**DOI:** 10.1101/2020.01.14.906982

**Authors:** Erika Rodríguez-Muñoz, Camilo Montes, Andrew J. Crawford

## Abstract

**Aim:** To evaluate the potential role of the orogeny of the Eastern Cordillera (EC) of the Colombian Andes and the Mérida Andes (MA) of Venezuela as drivers of vicariance between populations of 37 tetrapod lineages co-distributed on both flanks, through geological reconstruction and comparative phylogeographic analyses.

**Location:** Northwestern South America

**Methods:** We first reviewed and synthesized published geological data on the timing of uplift for the EC-MA. We then combined newly generated mitochondrial DNA sequence data with published datasets to create a comparative phylogeographic dataset for 37 independent tetrapod lineages. We reconstructed time-calibrated molecular phylogenies for each lineage under Bayesian inference to estimate divergence times between lineages located East and West of the Andes. We performed a comparative phylogeographic analysis of all lineages within each class of tetrapod using hierarchical approximate Bayesian computation (hABC) to test for synchronous vicariance across the EC-MA. To evaluate the potential role of life history in explaining variation in divergence times among lineages, we evaluated 13 general linear models (GLM) containing up to six variables each (maximum elevation, range size, body length, thermoregulation, type of dispersal, and taxonomic class).

**Results:** Our synthesis of geological evidence suggested that the EC-MA reached significant heights by 38–33 million years ago (Ma) along most of its length, and we reject the oft-cited date of 2–5 Ma. Based on mtDNA divergence from 37 lineages, however, the median estimated divergence time across the EC-MA was 3.26 Ma (SE = 2.84) in amphibians, 2.58 Ma (SE = 1.81) in birds, 2.99 Ma (SE = 4.68) in reptiles and 1.43 Ma (SE = 1.23) in mammals. Using Bayes Factors, the hypothesis for a single temporal divergence interval containing synchronous divergence events was supported for mammals and but not supported for amphibians, non-avian reptiles, or birds. Among the six life-history variables tested, only thermoregulation successfully explained variation in divergence times (minimum AICc, *R*^2^ = 0.10), with homeotherms showing more recent divergence relative to poikilotherms.

**Main conclusions:** Our results reject the hypothesis of the rise Andean Cordillera as driver of vicariance of lowland population because divergence dates are too recent and too asynchronous. We discuss alternative explanations, including dispersal through mountain passes, and suggest that changes in the climatic conditions during the Pliocene and Pleistocene interacted with tetrapod physiology, promoting older divergences in amphibians and reptiles relative to mammals and birds on an already established orogen.

## Introduction

Vicariance is a biogeographical model that describes the division of a widespread ancestral population into two daughter populations through the appearance of a barrier to gene flow that eventually leads to allopatric speciation (Crisci, 2001). Traditionally, such hypothesized barriers arise through geological changes, such as mountain uplift or the appearance of rivers. Environmental changes, such as aridification, can also promote spatial isolation of a formerly continuous population or meta-population system (Lomolino, Riddle & Whittaker, 2017). Most older methods of biogeographic analysis are based on a strictly vicariance model because dispersal was regarded as difficult to falsify (Rosen, 1978; Morrone & Crisci, 1995; Ronquist, 1997). Vicariance is also the most general and easiest to satisfy model of speciation, requiring only physical separation and the accumulation of nucleotide substitutions over time to create two daughter species from a common ancestor (Mayr, 1963; Wiens, 2004).

Despite the potential ubiquity of vicariance in promoting diversification of evolutionary lineages, many appreciable barriers, such as tall mountain ranges, appear to have conspecific populations on both sides. The effectiveness of a potential montane barrier may depend on the steepness of the environmental gradient and not on elevation *per se* (Janzen, 1967), suggesting that environmental heterogeneity might limit dispersal just as much, or perhaps more so, than simple physical obstruction. Therefore, their effectiveness in separating populations may have more to do with the interaction between environmental heterogeneity and organismal life history (Paz *et al*., 2015). For example, homeothermic animals likely have a higher tolerance to temperature heterogeneity compared to poikilotherms (Porter & Gates, 1969; Ghalambor, 2006). Life history variables could thus explain why the same potential barrier could differentially impact even closely related species.

We evaluate the vicariant model by looking across 37 lineages of tetrapods distributed on either side of the Eastern Cordillera of the northern Andean mountains of South America, specifically Colombia and Venezuela. The Andean Cordillera is the most extensive mountain range in the world, and the highest in America, stretching 9,000 km along the western coast of South America. Tropical mountains should represent an even stronger barrier to the local fauna and flora than temperate mountains (Janzen 1967; Ghalambor, 2006). Because the temperature in the tropical lowlands is relatively homogeneous across the landscape, species are expected to evolve narrower temperature tolerances compared to species that occupy high elevation zones and experience higher daily variations in temperature (McCain, 2009).

### Geology of the Andes

At its northern end, the Andes splits into three chains: The Western Cordillera (WC), the Central Cordillera (CC), and the Eastern Cordillera (EC) (Fig. 1). The EC extends 750 km from southern Colombia to northwestern Venezuela. The mean height of the EC is close to ~3,000 m.a.s.l. with summits reaching 5,500 m.a.s.l. The Mérida Andes (MA) extends 350 km from the northwestern Colombian-Venezuelan border (Fig. 1) and reaches a maximum elevation of ~5000 m.a.s.l.

**Figure 1.**
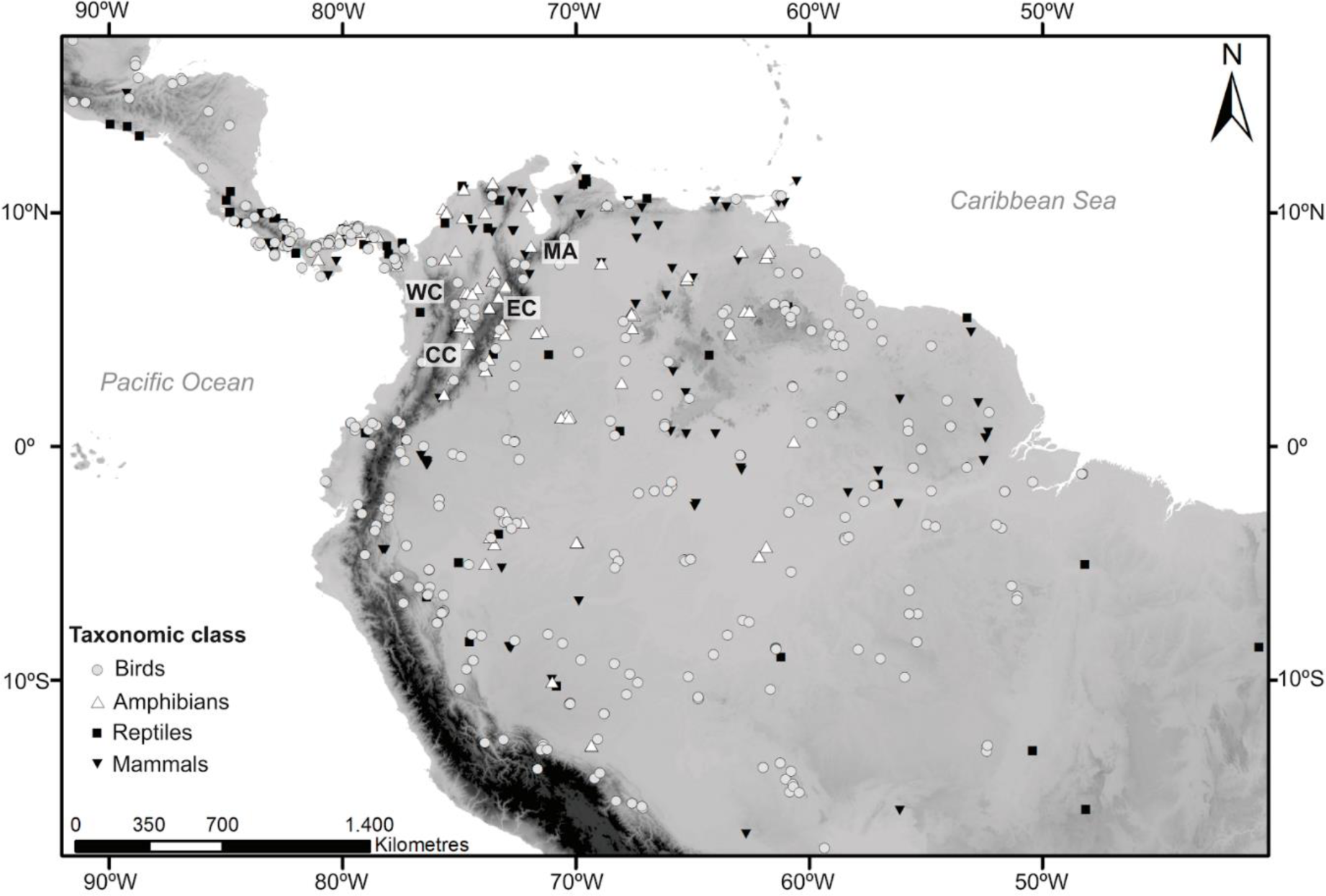
Sampling across in South America and Middle America per each of the four taxonomic classes of tetrapod studied here. The divisions of the northern Andes are labeled as Western Cordillera (WC), Central Cordillera (CC), Eastern Cordillera (EC), and the Mérida Andes as (MA).

During Cretaceous times most of northwestern South America was covered by shallow seas, the kind found on today’s continental platforms, perhaps only ~200m deep. Large, westward-flowing river systems, predating the Amazonian/Orinoco drainages dumped massive, thick sandstone strata along the western margin of the stable Guiana craton, today exposed, among others, in the Cocuy mountains of Colombia (Fabre, 1985). The record of these shallow marine conditions consists of limestone and fine-grained sedimentary rocks such as shale that generally grade eastward to coarser material such as sandstone and siltstone (Villamil & Arango, 1998). This record is preserved throughout the region in strata of that age, from the MA in the North, the Guajira Peninsula, the Santa Marta massif, the Perijá range, the CC, the Maracaibo block and the EC (see review in Sarmiento-Rojas et al., 2018). This Cretaceous shallow marine sequence (a passive margin) constitutes the base level from which to study the influence of elevation, the development of terrestrial barriers, and the effects of a dynamic landscape on biotas and diversification.

Arrival of oceanic-borne terranes to the Cretaceous passive margin of northwestern South America took place during one or several arc-continent collisions (see review in Montes et al., 2019). Collisions built relief along the margin of northwestern South America (see review in Bayona, 2018), shedding coarse-grained clastic material, sand and gravel, marking the start of a regional marine regression and the establishment of swampy conditions where shallow marine conditions had been prevalent for a long time. From this time onwards (latest Cretaceous to the south, to middle Paleocene in the north) we can define a primordial, probably discontinuous, Central Cordillera from Guayaquil to the Guajira peninsula. This orogen may have taken ~10 million years to propagate from south to north (see review in Montes et al., 2019). This range would have spawned the first Amazonian/Orinoco-style drainages, shutting down most or all westward-directed drainages of the region (Hoorn et al., 2010). We can therefore roughly reconstruct a north-south, single-cordillera configuration during latest Cretaceous to middle Eocene times in northwestern South America.

By late Eocene times, ~38–33 Ma, the geodynamic configuration of the northern Andes (Fig. 2A) changes and deformation starts to propagate eastwards into the domains of today’s EC (Mora et al., 2006; Ochoa et al., 2012; Bayona et al., 2013; Lamus et al., 2013). From this time onwards, we can document a young Magdalena Valley north of ~ 4°N, defined by a topographic depression between two flanking, linear, mostly continuous ranges (Horton et al., 2015). These ranges were high enough to provide potential energy for the erosion and transport of thick, coarse-grained deposits now preserved along both margins of the Magdalena Valley. South of ~4°N, the mountainous terrain of the young EC would have eased into the lowlands of Amazonia/Orinoco. Later, by middle Miocene times (Fig. 2B, ~15–13 Ma) deformation on the western flank of the EC cordillera propagates westward, building the flanks of the orogen (Restrepo-Pace et al., 2004). This process propagates southward, where middle Miocene strata gets involved in the deformation (Saeid et al., 2017). The Magdalena Valley is thus the result of a tectonic, not erosional, processes that modified an originally low-lying topography. Today, the Valley floor is flat, wide, and at the upper Valley, near 500 m.a.s.l., it defines the western edge of the EC.

**Figure 2A.**
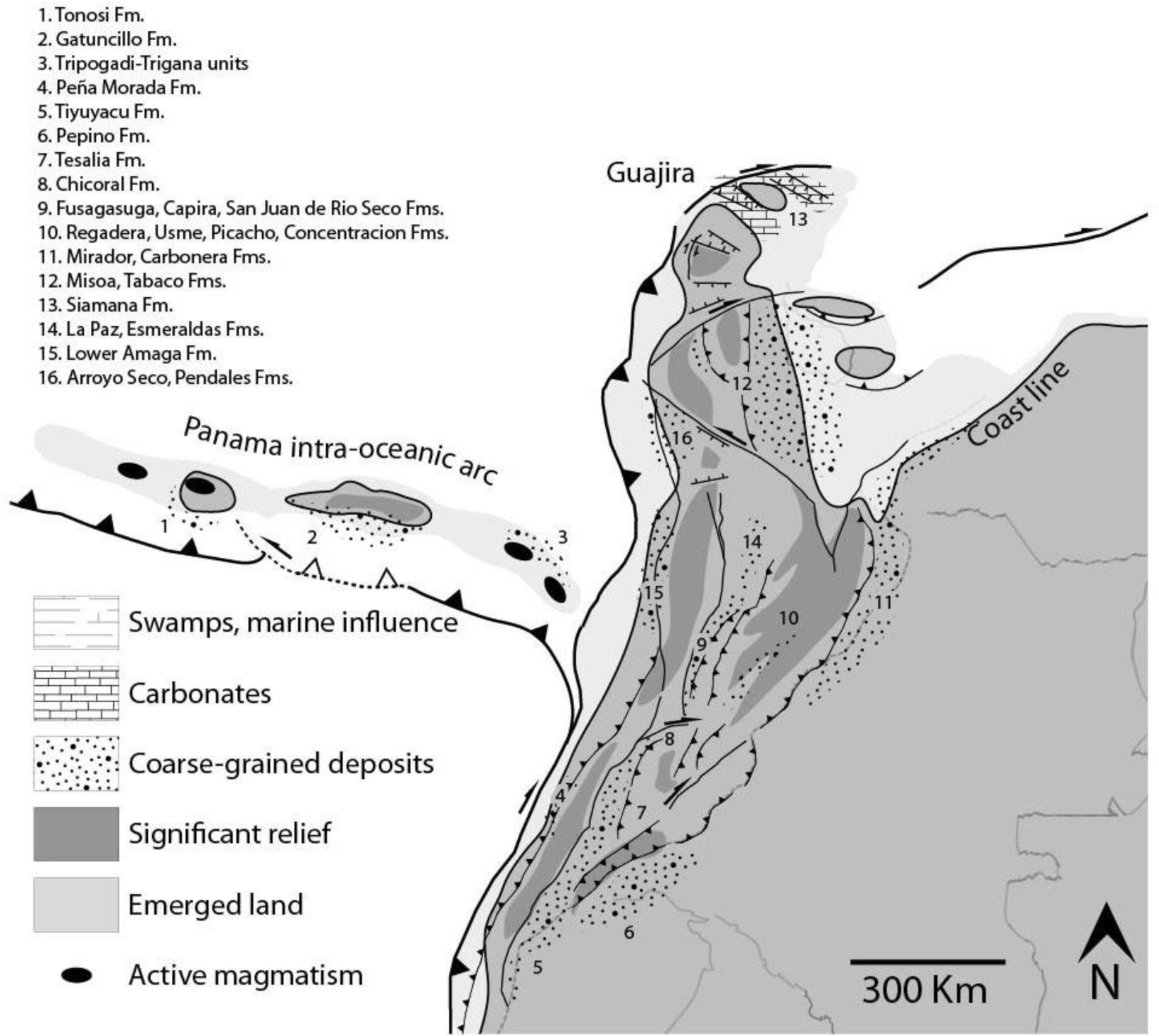
Palinspastic reconstruction of the northern Andes and southern Caribbean for latest Eocene-early Oligocene times (Montes et al., 2019). Political boundaries and the outline of Maracaibo Lake for reference. Note that the coarse-grained deposits (conglomerate and sandstone), common at this time, are used to track the presence and approximate location of relief, but not its magnitude (lithostratigraphic units after Bayona et al., 2008; Borrero et al., 2012; Caicedo and Roncancio, 1994; Cardona et al., 2012; Cardona et al., 2014; Christophoul et al., 2002; Gomez et al., 2005; Grosse, 1926; Herrera et al., 2012; Kolarsky et al., 1995; Moreno et al., 2015; Ochoa et al., 2012; Parnaud et al., 1995; Rodriguez and Sierra, 2010; Woodring, 1957).

**Figure 2B.**
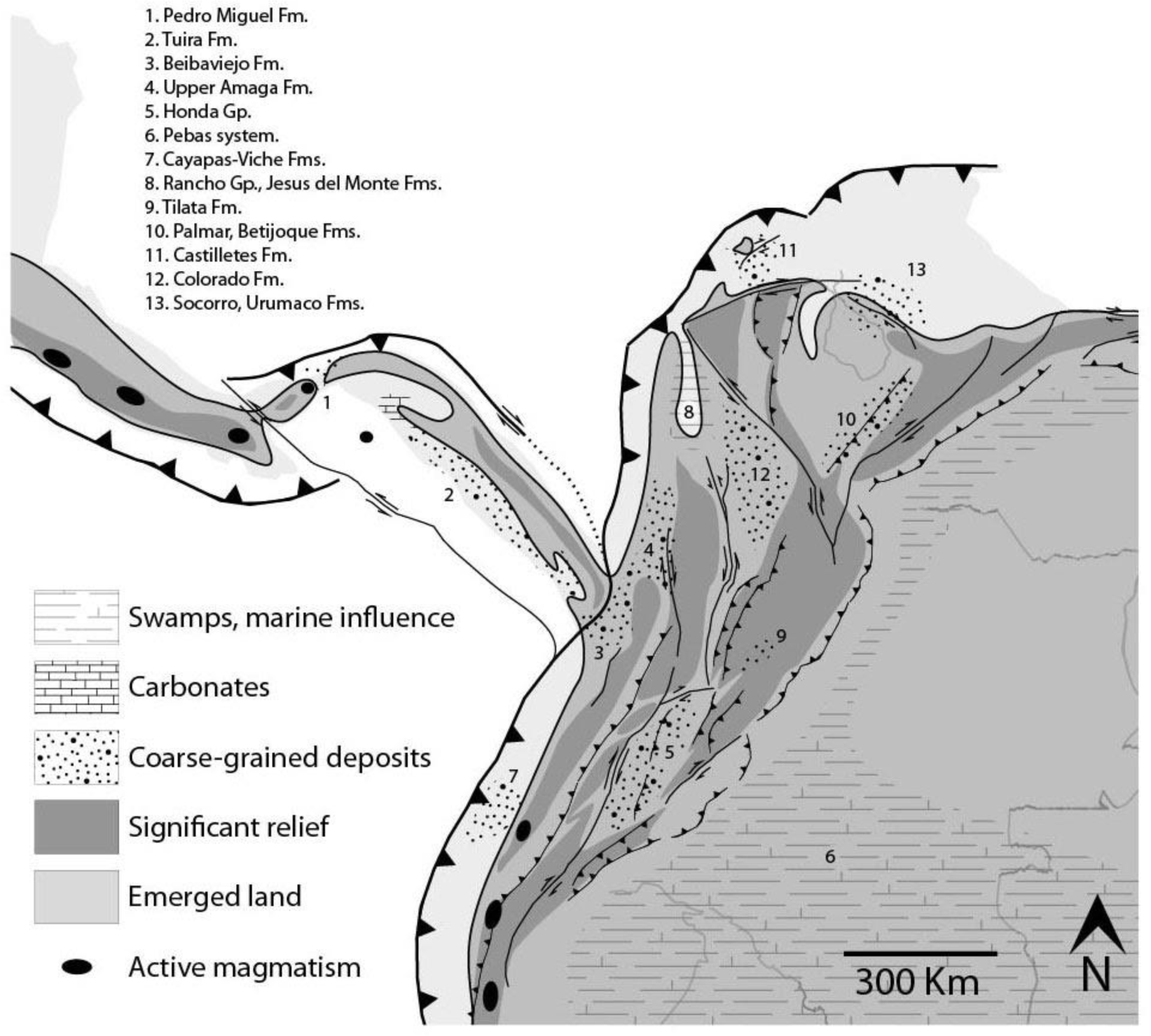
Palinspastic reconstruction of the northern Andes and southern Caribbean for for middle Miocene times (Montes et al., 2019). Political boundaries and the outline of Maracaibo Lake for reference. Most coarse-grained deposits at this time are sandy, from mostly fluvial and near-shore environments, and mark the segmentation of basins by rising mountain belts (Anderson et al., 2015; Barat et al., 2014; Borrero et al., 2012; Erikson et al., 2012; Farris et al., 2017; Gomez et al., 2005; Grosse, 1926; Guerrero, 1997; Hoorn et al., 2010; Leon et al., 2018; Montes et al., 2010; Moreno et al., 2015; Parnaud et al., 1995; Quiroz et al., 2010). Only two possible lowland passages are allowed at this time by these lithostratigraphic constraints and fish faunas: The Putumayo (Lundberg and Chernoff, 1992) to the South, and the Táchira corridor to the North.

The often-cited review paper about EC uplift (Gregory-Wodzicki, 2000), based on a classical paleobotanical study of the Bogotá plateau by van der Hammen et al. (1973), suggest that the modern elevation of the EC-MA was reached between 2–5 Ma. Such a date has commonly been taken as an absolute value representing simultaneous uplift along the entire length of the EC. Recent biomarker studies suggest that cooling revealed by paleobotanical studies (~9–12°C), may need to be revised downwards (3±1°C, Anderson et al., 2015). If that is the case, cooling may simply be the result of changing regional climates in the Miocene-Pliocene transition with no need for further surface uplift at all (Perez-Angel et al., 2017). Evidence for Andean uplift reviewed above suggests that its elevation history is much longer and more complex as revealed by coarse-grained strata on the flanks of the Andean ranges, and consistent with the recent biomarker studies.

In summary, the above synthesis of geological data, specifically of coarse-grained deposits, suggests that surface uplift of the Eastern Cordillera was already well underway in Eocene times (Fig. 2A), and nearly complete in middle Miocene times (Fig. 2B). Nonetheless, fossil fishes of Amazonian affinities found in the southern Magdalena Valley (the giant Pirarucú, *Arapaima gigas*, and others), require Magdalena-Amazonia biotic exchange until middle Miocene times (11 Ma; Lundberg and Chernoff, 1992). Such a pass may have existed where the Central Cordillera meets the southern end of the EC. Similarly, fossil catfish faunas of the upper Miocene Urumaco Formation in northwestern Falcon, Venezuela (Diaz de Gamero, 1996; Aguilera et al., 2013), and the Pliocene Castilletes Formation in the Guajira peninsula (Aguilera et al., 2013) suggest a lowland connection between Amazonia and the Caribbean coast at western end of the MA (Venezuela) as recently as ~ 3 Ma.

### Surface uplift and speciation

Andean uplift triggered changes in climatic and hydrologic conditions that led to the formation of the Amazon River system ~10.5 Ma (Figueiredo et al., 2009). This uplift is widely cited as having caused the primary divergence between ancestral populations on either side of the Andes. The rise of the EC, for example, is thought to have mediated the initial split between eastern and western populations of woodcreepers (*Dendrocincla*; Weir & Price, 2011), and between populations of the cane toad, *Rhinella marina*, with its recently recognized sister species, *R. horribilis* (Slade & Moritz, 1998). However, these and similar phylogeographic studies make strong assumptions about the timing and location of uplift of the EC-MA. First, many biologists assume that the Andes, or even a small section of the Andes such as the EC-MA, rose up synchronously across its entire length, without allowing for spatial heterogeneity in the timing of uplift. Second, even if uplift were spatially synchronous, at what point during the process of uplift should we assume migration ended? In the case of the EC-MA should we assume the date of initial uplift date of roughly 38 Ma for the separation of an ancestral population, or perhaps the final uplift data of 2 Ma based on palynological records near Bogotá in the middle of the EC (Van der Hammen et al.,1973), or some date in between? With such a long time period between the start and end of uplift of the EC-MA, this process could explain nearly any divergence data the investigator might obtain from evolutionary genetic analysis of her or his taxon of interest. Furthermore, some conspecific populations separated by the EC are not reciprocally monophyletic, demonstrating either a young separation date or continued migration over the mountains (Miller et al., 2008). Third, even if uplift were synchronous, and even if the investigador could connect without error a certain uplift date to divergence dates based on genetic data, this date of separation between eastern and western populations might not be relevant to other organisms with contrasting life histories (see above). Thus, any evaluation of the role of the Andes in driving primary divergence, i.e., vicariance, requires recognizing explicitly that even a section of the Andes, such as the EC-MA, is comprised of historically independent geological elements, and it requires a widely comparative approach to capture the variation among lineages in their response to changes in the environmental conditions.

To evaluate the role of the uplift of the EC-MA mountain chains on genetic variation, we chose to work with tetrapods, as these are similarly large and terrestrial organisms for which ample phylogeographic data sets are available online. At the same time this clade is also diverse enough to capture remarkable variation in potentially relevant life-history and eco-physiological traits that allow us to evaluate their possible effect on divergence times across the EC-MA. We used comparative phylogeography and hierarchical approximate Bayesian computation (hACB) to test the following hypotheses, in order of complexity. H1) The separation of eastern and western clades within each species or genus was simultaneous among all taxa studied here, supporting the classic vicariance model and little role for life-history variation affecting the timing of divergence. This hypothesis predicts that A) our hABC analysis will reveal a single interval containing all divergence events, and B) according to off-cited estimates, the divergence time interval will be around 2–5 Ma, whereas our re-analysis of geological evidence (above) suggests much older dates. The next three hypotheses suggest that barriers are the result of organismal-environmental interactions. Thus, how the rise of the EC-MA affected each species depends on the eco-physiological traits of each species. H2) The separation of eastern and western clades was asynchronous among taxa and this variation can be explained by differences in elevation and geographic range. As the EC-MA rose up, lowland taxa would have separated first and organisms with wider distributional ranges, higher dispersal abilities, and/or broader tolerance of landscape heterogeneity should be less affected by the environmental gradient generated by mountain uplift. This hypothesis is consistent with a traditional vicariance model but allows for some variation in divergence times among co-distributed taxa. H2 thus predicts that species with higher elevation ranges and widespread species will show younger divergence times relative to lowland and narrowly distributed species since they have lower dispersal abilities and more specific habitat requirements. H3) The effectiveness of a montane barrier will be stronger for small-bodied species since they are poor dispersers compared to bigger species (Jenkins et al., 2007; Paz et al., 2015), predicting that divergences vary according to body size, with older divergences in smaller species. H4) Variation in divergence times (if any) can be explained by physiological traits, as follows. 4A) homeotherms will show shallower divergences across the Andes, while poikilotherms will show deeper divergences due to their susceptibility to environmental temperature variation, reducing their dispersal across the incipient environmental gradient created by mountain uplift (Porter & Gates, 1969; Buckley et al., 2012). 4B) Flying species (bats and birds) will show younger divergences respect to non-volant species since the former latter should disperse much more readily over the nascent Andes.

## Materials and methods

### Species selection and genetic data collection

To re-assess the role of the Eastern Cordillera and Mérida Andes, EC-MA, in separating ecologically diverse lowland taxa into eastern and western populations, we combined new information with published data sets representing clades with *cis-*and *trans*-Andean distributions, i.e., East and West of the EC-MA, respectively. We restricted the selection of ingroup samples according to the following filters. First, we included only tetrapods (see Introduction). Second, each set of related populations must comprise a monophyletic group, independently of species names currently assigned to the populations. Populations on either side of the EC-MA, however, were not required to be monophyletic to be included in phylogenetic analyses (but see below regarding assumptions of comparative phylogeographic analyses). Third, the chosen clade must include a maximum of one species on at least one side of the EC-MA, while the opposite side could contain the same species, a sister species, or a sister clade with up to a maximum of two named species (Fig. 3). In this way, we sought to limit our study to potential cases of primary vicariance across the mountains, while avoiding more ancient divergence events obscured by more recent speciation on either side. Fourth, since we were testing a model of vicariance of lowland taxa by mountain uplift, we excluded tetrapod species whose maximum elevation range exceeded 2,000 m.a.s.l. Fifth, in order to exploit coalescent-based analytical tools, we also required datasets to have a minimum of three conspecific individuals sampled on each side of the EC-MA. Sixth, because very few published studies passing the preceding filters also included nuclear DNA data, we were obliged to limit our comparative analyses to mitochondrial DNA (mtDNA) sequence data available in GenBank, in the Barcode of Life Database (BoLD; Ratnasingham & Hebert, 2007), or, in the case of frogs, using new data reported here for the first time. We found 30 published data sets meeting the above requirements, and to these we added five previously unpublished datasets from Colombian frogs. Among these 37 data sets, the total number of lineages per taxonomic class was nine amphibians, five from reptiles, 17 bird data sets, and six from mammals (Table 1). GenBank accession numbers are provided in Table S1 (Accession numbers for new data will be supplied prior to final acceptance of this manuscript for publication.)

**Figure 3.**
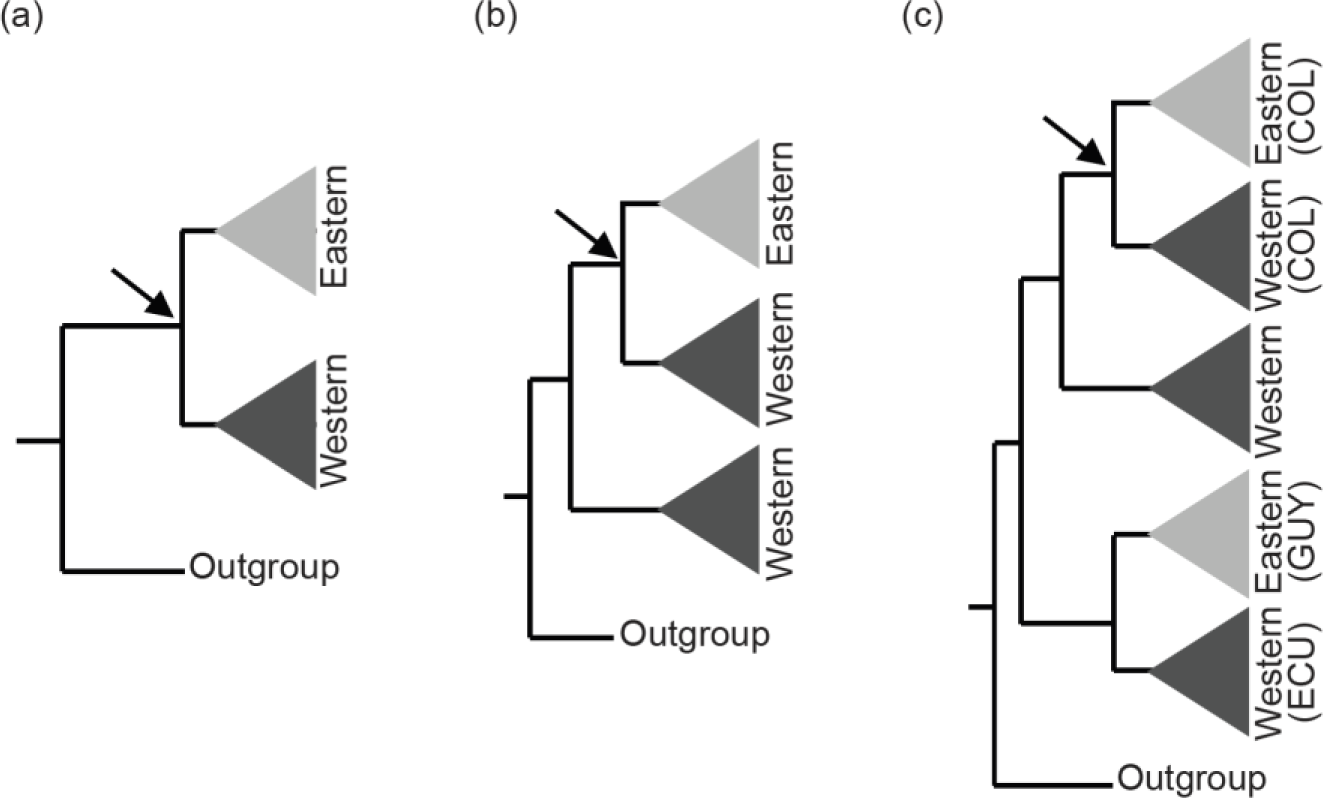
The assumptions of the comparative phylogeographic analysis in msBayes using hABC (see Methods) required sub-sampling the Bayesian phylogenetic consensus trees, as follows. Arrows mark the subclades selected for analysis by hABC according to the possible tree topology. For reciprocally monophyletic groups (a) all data were used. For paraphyletic groups (b) we sampled only the clade that fit the two-population model assumed by msBayes. For polyphyletic groups (c) we selected the clade that adjusted to the two-population model with sampling localities geographically closest to the Eastern Cordillera (EC). COL = Colombia, ECU = Ecuador, GUY = Guiana.

**Table 1.**
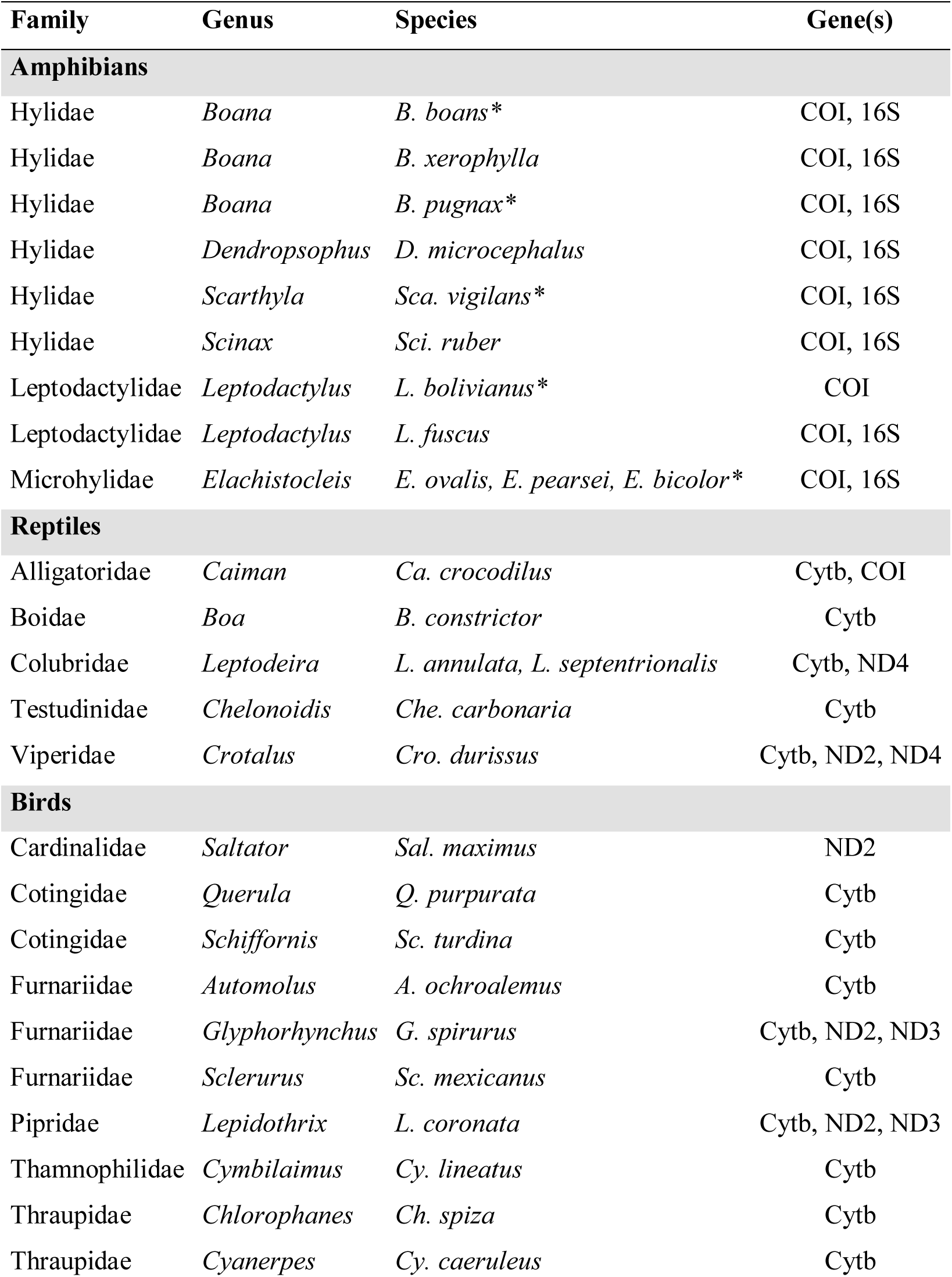
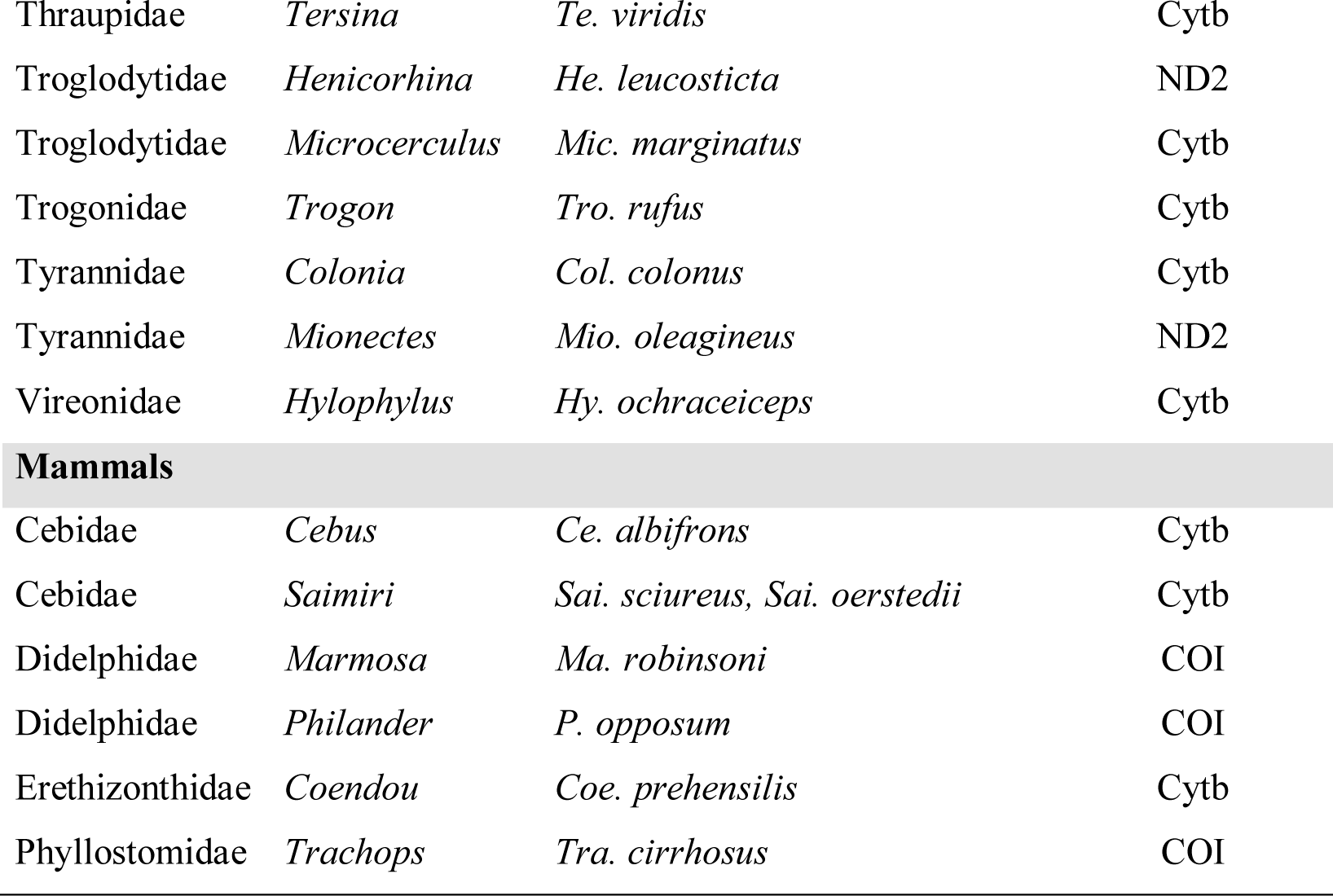
List of 37 taxa among four taxonomic classes of tetrapods analyzed in the present work, along with abbreviations for the mitochondrial genes used in phylogenetic and comparative phylogeographic analyses. Data published for the first time here are marked with an *. GenBank numbers for new and previously published data are provided in Supplementary Online Table S1

| Family | Genus | Species | Gene(s) |
| --- | --- | --- | --- |
| **Amphibians** |  |  |  |
| Hylidae | *Boana* | *B. boans* * | COI, 16S |
| Hylidae | *Boana* | *B. xerophylla* | COI, 16S |
| Hylidae | *Boana* | *B. pugnax* * | COI, 16S |
| Hylidae | *Dendropsophus* | *D. microcephalus* | COI, 16S |
| Hylidae | *Scarthyla* | *Sca. vigilans* * | COI, 16S |
| Hylidae | *Scinax* | *Sci. ruber* | COI, 16S |
| Leptodactylidae | *Leptodactylus* | *L. bolivianus* * | COI |
| Leptodactylidae | *Leptodactylus* | *L. fuscus* | COI, 16S |
| Microhylidae | *Elachistocleis* | *E. ovalis*, *E. pearsei*, *E. bicolor* * | COI, 16S |
| **Reptiles** |  |  |  |
| Alligatoridae | *Caiman* | *Ca. crocodilus* | Cytb, COI |
| Boidae | *Boa* | *B. constrictor* | Cytb |
| Colubridae | *Leptodeira* | *L. annulata*, *L. septentrionalis* | Cytb, ND4 |
| Testudinidae | *Chelonoidis* | *Che. carbonaria* | Cytb |
| Viperidae | *Crotalus* | *Cro. durissus* | Cytb, ND2, ND4 |
| **Birds** |  |  |  |
| Cardinalidae | *Saltator* | *Sal. maximus* | ND2 |
| Cotingidae | *Querula* | *Q. purpurata* | Cytb |
| Cotingidae | *Schiffornis* | *Sc. turdina* | Cytb |
| Furnariidae | *Automolus* | *A. ochroalemus* | Cytb |
| Furnariidae | *Glyphorhynchus* | *G. spirurus* | Cytb, ND2, ND3 |
| Furnariidae | *Sclerurus* | *Sc. mexicanus* | Cytb |
| Pipridae | *Lepidothrix* | *L. coronata* | Cytb, ND2, ND3 |
| Thamnophilidae | *Cymbilaimus* | *Cy. lineatus* | Cytb |
| Thraupidae | *Chlorophanes* | *Ch. spiza* | Cytb |
| Thraupidae | *Cyanerpes* | *Cy. caeruleus* | Cytb |
| Thraupidae | *Tersina* | *Te. viridis* | Cytb |
| Troglodytidae | *Henicorhina* | *He. leucosticta* | ND2 |
| Troglodytidae | *Microcerculus* | *Mic. marginatus* | Cytb |
| Trogonidae | *Trogon* | *Tro. rufus* | Cytb |
| Tyrannidae | *Colonia* | *Col. colonus* | Cytb |
| Tyrannidae | *Mionectes* | *Mio. oleagineus* | ND2 |
| Vireonidae | *Hylophylus* | *Hy. ochraceiceps* | Cytb |
| **Mammals** |  |  |  |
| Cebidae | *Cebus* | *Ce. albifrons* | Cytb |
| Cebidae | *Saimiri* | *Sai. sciureus, Sai. oerstedii* | Cytb |
| Didelphidae | *Marmosa* | *Ma. robinsoni* | COI |
| Didelphidae | *Philander* | *P. opossum* | COI |
| Erethizontidae | *Coendou* | *Coe. prehensilis* | Cytb |
| Phyllostomidae | *Trachops* | *Tra. cirrhosus* | COI |

### Molecular phylogenetic analyses

To the above sampling we added one or more outgroup samples to each dataset in order to root trees and to assist in temporal calibrations of molecular phylogenies. We searched the literature for time-calibrated trees containing the ingroup of interest, and we selected as outgroup the closest species that had published DNA sequences of the same gene or genes as available for the ingroup. When available, we added additional outgroup species from the same genus or family to reduce uncertainty on node ages. DNA sequence data sets were aligned independently for each genus and each gene using ClustalW v. 1.74 (Larkin et al., 2007) with gap opening costs set to 20 for protein-coding genes and 16 for ribosomal RNA genes. PartitionFinder2 (Lanfear *et al*., 2012) was applied independently to each alignment to select the best-fit partitioning schemes and models of nucleotide substitution. Potential partitions considered included by gene and by codon position.

To estimate divergence times independently for each lineage between eastern and western samples, we employed the Bayesian MCMC molecular phylogenetic software BEAST v. 2.4.7 (Bouckaert *et al*., 2014). We ran two chains for 80 million generations, sampled every 5,000 steps with a coalescent constant-size tree prior since each data set included population samples and it is the most suitable prior for describing relationships within and between populations. Searches started from a random tree and assumed an uncorrelated lognormal relaxed molecular clock. Datasets with multiple mitochondrial genes were concatenated under the assumption of no recombination in tetrapod mtDNA. For birds, we set the prior on the substitution rate using a lognormal distribution that included the well-established rate for an avian mtDNA molecular clock of 2% total divergence per million years (Weir & Schluter, 2008). For anuran mtDNA, we assumed a lognormally distributed substitution rate prior with a mean of 0.00955 divergence per lineage per million years and a range from 0.0074 to 0.01225, corresponding to the 2.5% and 97.5% quantiles, respectively, derived from the silent site divergence rates reported in Crawford (2003). For reptiles and mammals, we employed only secondary calibrations derived from published timetrees (Appendix S2), and we did not used substitution rates because the MCMC chains did not converged when we constrained used both the node ages priors and with substitution rates.

### Comparative phylogeographic analysis

We used the software MTML-msBayes (v. 20170511) (Overcast et al., 2017) to evaluate the degree of temporal coincidence of divergence events among *n* = 37 tetrapod taxa geographically divided by the EC-MA. MTML-msBayes uses hierarchical approximate Bayesian computation (hABC) to combine data from multiple co-distributed lineages into a global coalescent analysis that includes a hyperparameter (Ψ) describing the number of time intervals necessary to account for the observed range of divergence times among *n* taxa, with a single time interval (Ψ = 1) implying simultaneous divergence of all lineages, up to a maximum of Ψ = *n*, i.e., divergence across the EC-MA occured at a statistically distinct point in time for each taxon (Huang et al., 2011).

The first step in the hABC analysis is estimating population genetic summary statistics for each lineage. Subsequently, data sets of the same size as the observed data are simulated under a coalescent model using parameter values drawn from a prior distribution, and summary statistics are estimated from each simulated dataset. Finally, an acceptance/rejection algorithm is applied to obtain a sample from the posterior distribution by comparing the summary statistics of each simulated dataset with those from the observed data (Huang et al., 2011). As suggested by Hickerson *et al*. (2007) for small datasets, we employed the genetic distance between populations, π_b_ (Charlesworth, 1998), as the summary statistic for the estimation of the parameters Ψ, the mean divergence time E(τ), where τ is the age of the splitting event that divided the ancestral population, and for Ω, the dispersion index of τ [var(τ)/E(τ)]. Ω < 0.0 is commonly used as criterion for simultaneous divergence for multiple pairs of codistributed lineages (Hickerson et al., 2007). We assumed that mitochondrial genes are in perfect linkage disequilibrium, and therefore treated multi-gene datasets as single, concatenated sequences in hABC, and excluded any sample missing data from two or more genes. MTML-msBayes assumes a two-population model (Fig. 3a) that is violated by deep phylogeographic structure on either side of the proposed barrier (Huang et al., 2011). Therefore, we checked tree topologies obtained using BEAST (see above) and for those few taxa with multiple lineages on one side of the EC-MA we included only samples from the clade that best fit the two-population model implemented in MTML-msBayes. Thus, for paraphyletic and polyphyletic groups, we included the clade with the most samples (to obtain better estimates of population genetic parameters) and/or clades geographically positioned most closely to the EC-MA (Fig. 3c).

The number of possible divergence models increases when more taxa are added, thus the computer will not be able to evaluate all the possible models properly when implementing a single analysis with the 37 lineages (Oaks et al., 2013), hence we ran one independent analysis for each of the four classes of tetrapods. We set the upper limit of τ to the oldest mean split age for each class according to our BEAST analysis. Divergence times estimated from MTML-msBayes are in coalescent units, thus the conversion to millions of years assumed roughly equal sex ratios, haploid and maternally inherited mtDNA, and was made following the equation *t* = τ*θ*_Ave_/*μ*, where *θ*_Ave_ is the average effective population size for each taxonomic class estimated by MTML-msBayes and *μ* is the neutral mutation rate per site per generation. Values of *μ* for frogs were taken from Crawford (2003) and for birds we followed Dolman & Joseph (2011). For mammals and non-avian reptiles, we used specific rates per genus (see Appendix S3). Further details regarding estimating coalescent-based divergence times, hyperpriors, substitution rates, and generation times assumed for each taxon are given in Appendix S3. Hyper-posteriors were estimated from 1,000 accepted draws from 1.5 million simulations. We made a local linear regression of the accepted parameter values obtained by the acceptance/rejection step in order to improve the posterior estimation. We used Bayes factors (BF; Kass & Raftery, 1995) to evaluate the relative posterior support for the number of divergence pulses. To estimate the timing of each divergence interval and the species contained in each, we constrained Ψ to the value with maximum BF and repeated the analysis as outlined above (Paz et al. 2015).

### Life-history determinants of divergence times

We selected six life-history variables potentially related to dispersal abilities that could influence divergence times: body size as length, geographic range, upper elevation limit, type of locomotion (flying versus not flying), thermoregulation (homeotherm versus poikilotherm) and taxonomic class (amphibians, non-avian reptiles, birds, mammals). We obtained snout-vent length of amphibians from the AmphiBIO database (Oliveira et al., 2017). For reptiles we obtained body-size data from the literature (Savage, 2002; Bartlett & Bartlett, 2003; Böhm et al., 2013; Fowler, 2018). We obtained body-size data for birds from the Handbook of the Birds of the World (del Hoyo et al., 2018), and for mammals we consulted the amniote life-history database (Myhrvold et al., 2015). We used species distribution shapefiles from the IUCN Red List of Threatened Species (2018) to estimate the geographic ranges of species in km^2^. We obtained upper elevational-limit data from the IUCN Red List of Threatened Species (2018), Amphibian Species of the World (Frost, 2018), and the Handbook of the Birds of the World. The variables mean divergence time and body size were transformed with the natural logarithm function to better meet assumptions of normality and homoscedasticity. Based on the above six life-history variables we defined a total of 13 *a priori* general linear models (GLM, Table 2) that could potentially explain variation in divergence times. We performed an AICc model selection procedure using R v. 3.5.1 (R Core Team, 2018) to determine which variables are more relevant in determining divergence times.

**Table 2.** List of the thirteen models selected *a priori* that could potentially explain differences in divergence times among data sets, the biological relevance of each model and their corrected Akaike Information Criterion (AICc) coefficients. Linear combination of variables is represented by ‘+’ symbol, while interaction between variables is represented by ‘×’.

| Model | Biological relevance | AICc | ΔAICc |
| --- | --- | --- | --- |
| Temperature regulation | Homeotherms can face environmental gradients, facilitating dispersal. | 86.79 | 0 |
| Upper elevation limit | Physical barriers would have greater impact for species with lower elevation limits. | 89.71 | 2.92 |
| Class | Differences in divergence times rely only in taxonomy. | 89.94 | 3.15 |
| Upper elevation limit × Temperature regulation | Homeotherms can face lower temperatures associated with elevation gradients and thus could reach higher elevations. | 90.61 | 3.81 |
| Body size | It is easier for larger organisms to disperse. | 91.07 | 4.28 |
| Dispersal | Flying organisms would be better dispersers. | 91.3 | 4.51 |
| Geographic range | Organisms with wider geographic ranges can be considered generalists and thus better dispersers. | 91.34 | 4.55 |
| Geographic range + Upper elevation limit | Species with wide geographic and elevation ranges. | 91.45 | 4.66 |
| Body size + Geographic range | Large and generalist species could disperse wider distances. | 93.15 | 6.36 |
| Body size + Geographic range + Upper elevation limit | Large species with wide geographic ranges and higher elevation limits would be more tolerant to environmental heterogeneity and would be more able to disperse more. | 93.33 | 6.54 |
| Body size × Class | Body size can vary across tetrapod classes,<br>influencing dispersal abilities. | 96.65 | 9.86 |
| Geographic range ×<br>Class | Geographic range could vary depending on the class<br>the organisms belong. | 97.97 | 11.18 |
| Upper elevation limit ×<br>Class | Elevation limit could differ among classes. | 98.24 | 11.45 |

## Results

### Divergence time estimation and hABC

We defined our nodes of interest as the youngest node that had descendents on both sides of the EC-MA. For frogs, the age of the node that split east and west populations in BEAST analysis ranged from 1.09 to 10.3 Ma (Fig. 4a), for birds 0.72 to 8.04 Ma (Fig. 4b), from 0.84 to 12.62 Ma in reptiles (Fig. 4c), and from 0.36 to 3.69 Ma among lineages of mammals (Fig. 4d). The comparison of posterior distributions supported strongly a scenario of asynchronous divergence in amphibians (BF = 0), reptiles (BF = 0) and birds (BF = 0), while for mammals the scenario of simultaneous divergence was moderately supported (BF = 3.29). For amphibians, BF values provided support for four divergence interval among the 9 lineages (BF = 4.12; Appendix S4, Fig. S1a), three pulses among the 5 reptile lineages (BF = 2.58; Appendix S4, Fig. S1b), and six among the 17 avian lineages (BF = 5.63; Appendix S4, Fig. S1c).

**Figure 4.**
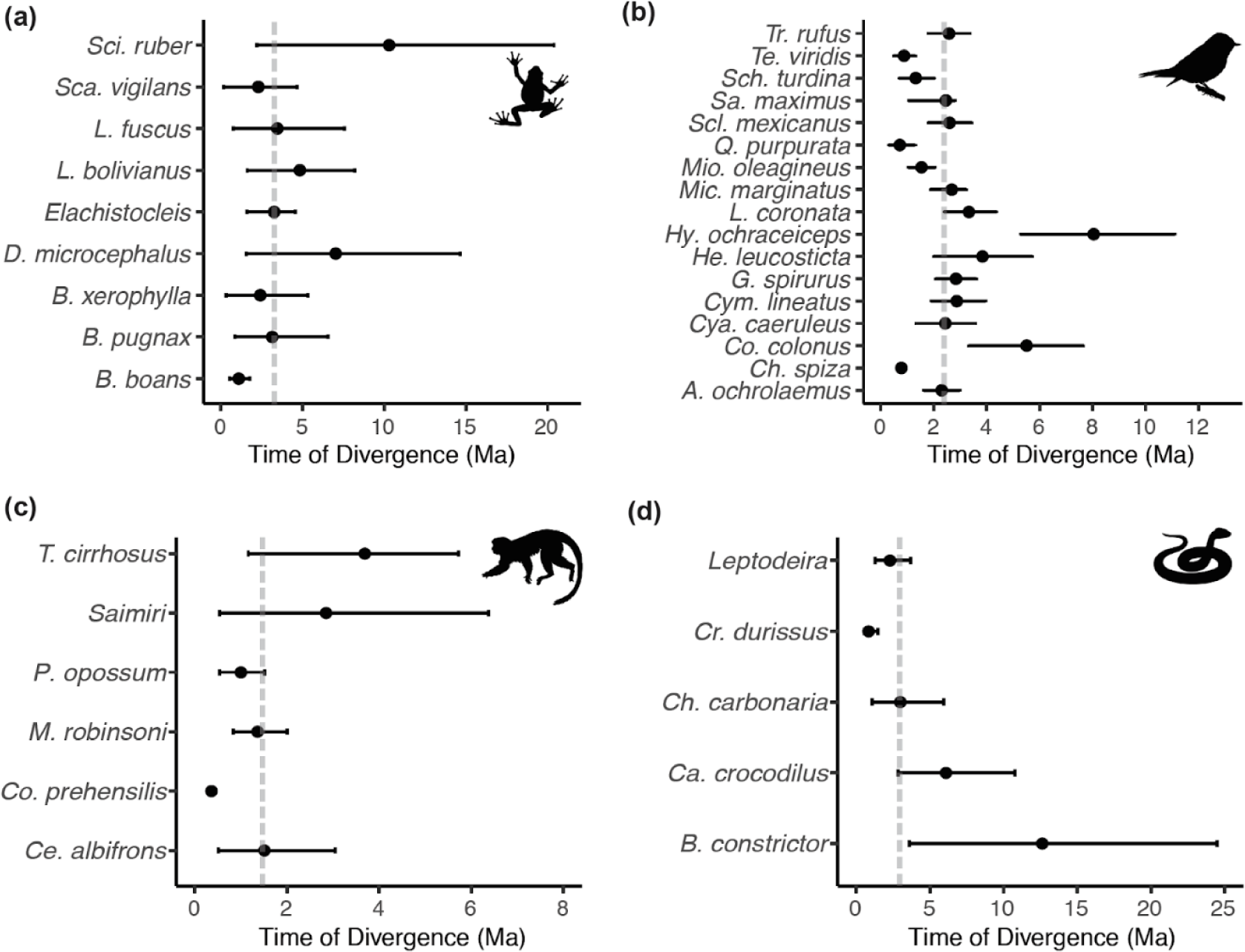
Distribution of divergence time estimates between eastern and western lineages of (a) amphibians, (b) birds, (c) mammals and (d) non-avian reptiles, estimated by Bayesian MCMC relaxed-clock phylogenetic analysis of mitochondrial DNA. Dots indicate the mean divergence time and dotted lines indicate the median divergence time within each class. Median divergence time in amphibians was 3.14 million years ago (Ma), 2.99 Ma in non-avian reptiles, 2.58 Ma in birds, and 1.03 Ma in mammals.

The timing of intervals of divergence events estimated by coalescent analysis and the participating species within each interval are given in Table 3. For each taxon, the divergence times across the EC-MA estimated from phylogenetic analyses largely agreed with the coalescent-based results. We performed a sign test within each taxonomic class to evaluate differences in divergence times estimated by the Bayesian phylogenetic approach (BEAST) versus the explicitly coalescent approach (MTML-msbayes). Only birds showed a significant difference, where the divergence time estimates using hABC were older than those estimated using BEAST (n = 17, *p* = 0.00235; Appendix S4), for the other the other taxa, we did not find significant differences (Amphibians: n = 9, *p* = 0.1797; Reptiles: n = 5, *p* = 1; Mammals: n = 6, *p* = 0.0625; Appendix S4).

**Table 3.** Number, grouping, and timing of each divergence interval (Ψ) estimated by hierarchical approximate Bayesian computation (hABC) as implemented in the software, MTML-msBayes. Intervals are independently estimated within each of the four taxonomic classes. Complete generic level names are found in Table 1. Ma = millions of years ago. CI = posterior credibility interval.

| Number of events ($\Psi$) | Set of species | Time (Ma) | 95% CI |
| --- | --- | --- | --- |
| **Amphibians** |  |  |  |
| $\Psi_1$ | *Boana boans*, *Sca. vigilans* | 0.13 | 0 – 0.45 |
| $\Psi_2$ | *Boana xerophylla*, *Elachistocleis*, *Leptodactylus fuscus* | 2.14 | 0.85 – 3.5 |
| $\Psi_3$ | *Boana pugnax*, *Den. microcephalus* | 3.62 | 1.97 – 5.83 |
| $\Psi_4$ | *Leptodactylus bolivianus*, *Sci. ruber* | 7.57 | 6.29 – 8.69 |
| **Reptiles** |  |  |  |
| $\Psi_1$ | *Cr. durissus*, *Che. carbonaria* | 1.9 | 0.51 – 3.02 |
| $\Psi_2$ | *Caiman crocodilus* | 4.59 | 1.5 – 8.66 |
| $\Psi_3$ | *Boa constrictor*, *Leptodeira* | 9.05 | 6.3 – 11.8 |
| **Birds** |  |  |  |
| $\Psi_1$ | *Chl. spiza*, *Q. purpurata*, *Scl. mexicanus*, *Te. viridis* | 0.23 | 0 – 0.47 |
| $\Psi_2$ | *Cya. caeruleus*, *Cym. lineatus*, *Mio. oleagineus* | 0.65 | 0 – 1.2 |
| $\Psi_3$ | *Col. colonus*, *Le. coronata*, *Sa. maximus* | 1.53 | 0.65 – 2.33 |
| $\Psi_4$ | *G. spirurus*, *He. leucosticta*, *Sch. turdina* | 2.10 | 1.22 – 2.96 |
| $\Psi_5$ | *A. ochroalemus*, *Mic. marginatus*, *Tro. rufus* | 2.67 | 1.62 – 4.06 |
| $\Psi_6$ | *Hy. ochraceiceps* | 7.73 | 6.07 – 9.44 |
| **Mammals** |  |  |  |
| $\Psi_1$ | *Ce. albifrons*, *Co. prehensilis*, *Ma. robinsoni*, *P. opossum*, *Saimiri*, *Tra. cirrhosus* | 0.36 | 0.31 – 0.43 |

### Life-history determinants of divergence times

Among the thirteen models we evaluated to explain variation in divergence times, the best-fit model contained only a single eco-physiological variable, ‘thermoregulation’ (AICc = 91.28; Table 2), with homeothermic species showing younger divergence times relative to poikilotherms (Fig. 5). However, this model explained only a small portion of the variation (*R^2^* = 0.10, Adjusted *R^2^* = 0.08). The second best-fitting model explained a little more of the variation and contained only the variable, ‘taxonomic class’ (ΔAICc = 1.49, *R^2^* = 0.19, Adjusted *R^2^* = 0.11; Table 2). Body size and geographic range appear to be the least informative predictors of divergence across the EC-MA since models including these variables (single variable, within linear combinations, or as interaction terms) have the highest AICc values.

**Figure 5.**
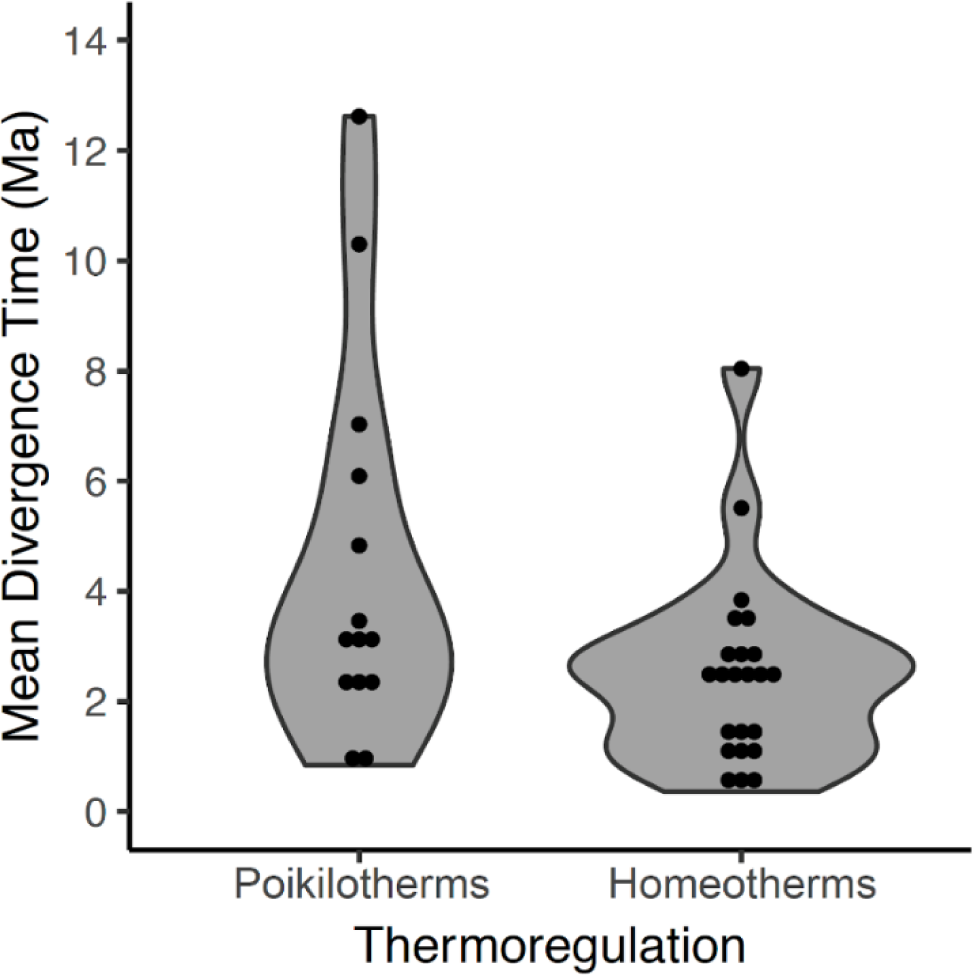
Mean divergence times (in million years, Ma) of eastern and western populations as estimated by Bayesian inference for homeotherm and poikilotherm tetrapods. The points represent the mean divergence time of each of 37 data sets.

## Discussion

### Pulses of divergence and Andean uplift

Comparison of phylogeographic patterns among co-distributed lineages allows the researcher to test geological events and landscape features as possible determinants of speciation that shaped current geographic patterns of biodiversity (Hickerson et al., 2010). The typical null hypothesis assumes that co-distributed species share a common biogeographic history influenced by the same historical events (Arbogast & Kenagy, 2008). Our data are not consistent with this hypothesis, and our results demonstrate the occurrence of several divergence pulses where the estimated divergence times, τ, vary so widely among taxa within each of the four taxonomic classes of tetrapod that no single time interval could account for this variation. The amphibian, non-avian reptile, and bird data sets each have at least one divergence pulse in which the mean τ is older or younger than the traditional dating of EC-MA uplift (2–5 Ma; Table 3). Only in mammals did the hABC framework support a single divergence time interval. However, at 0.31–0.43 Ma, this divergence time interval was far younger than the traditionally proposed divergence time across the EC-MA (Table 3). Thus, the variation in divergence times among taxa, as well as the number of splits outside the traditional 2–5 Ma interval, suggest we need to reconsider our evolutionary and geological models of the uplift of the EC-MA mountain chains.

### Differences in phylogeographic approaches

Birds showed a significantly older divergence time estimates using hABC relative to BEAST (Appendix S4, Table S2), which is unexpected since the divergence of an ancestral population should occur posterior to gene divergence, assuming no post-divergence migration (Nei & Li, 1979). This discrepancy could be attributed to the models used by each software, where BEAST estimates a phylogenetic timetree assuming complex models of substitution while MTML-msBayes uses a coalescent approach. However, this explanation would predict similar discrepancies in estimate divergence times across all 4 taxonomic classes, which was not the case. Avian taxa, however, showed substantial polyphyly with respect to the two sides of the EC, which may explain why mean gene divergence across the EC could be younger than the population divergence.

We employed mtDNA to infer phylogenetic relationships and the number and the timing of divergence events. This marker is widely used in comparative phylogeographic studies for several reasons. Its high mutation rate and lack of recombination provide ample and detailed information needed to reconstruct the genealogy of samples with more precision that found for any other single marker. The fact that its haploid and maternally inherited means genetic drift is at least 4-fold faster, which greatly lowers the probability of incomplete lineage sorting (ILS; Avise et al., 1987; Hudson & Coyne, 2002). Finally, the conserved structure of the mitochondrial genome makes it the best single marker for comparative analyses across diverse animal species (Carnaval et al., 2009), though mtDNA does not always predict variation in nuclear DNA (nDNA; Toews & Brelsford, 2012). In some cases, discordances between mtDNA and nDNA phylogenies are observed, so analysis including different loci are recommended to confirm phylogeographic histories. In this case in particular, we could not find DNA samples from multiple loci that could constitute a representative geographic sampling that could help to answer our research questions. Coalescent inference based on single-locus information could be inaccurate (Edwards et al., 2000). However, the approach used in hABC takes into account the intrinsic stochasticity in single-locus coalescence genealogies across different taxa and moreover, estimation of population divergence time (τ) does not improve considerably until eight nuclear loci are incorporated (Huang et al., 2011).

### Ecological factors and divergence times

The only life-history variable that was found to be associated with divergence times was thermal physiology, such that poikilotherms had older divergences across the EC-MA than homeotherms. Poikilotherm dispersal is restricted by temperature and these animals cannot perform across as wide a range of environmental temperatures as homeotherms can (Porter & Gates, 1969). Tropical lowland poikilotherms have restricted altitudinal ranges, narrow temperature tolerances and limited acclimation responses compared to high elevation and temperate poikilotherms species (Ghalambor, 2006).

These physiological trends are reflected in variation in divergence times among South American tetrapods, such that amphibians and non-avian reptiles tend to show Miocene to Pliocene intraspecific divergences, whereas mammals and bird species tend to show Late Pliocene to Pleistocene divergences (Turchetto-Zolet et al., 2013). Temperature oscillations may determine the altitudinal extent of an ecosystem. Quaternary glaciation events starting 2.58 Ma compressed vegetation belts downwards, while in the interglacial cycles these belts expanded and reached higher elevations (Hooghiemstra & Van der Hammen, 2004; Ramírez-Barahona & Eguiarte, 2013). As we argue in more detail below, interglacial episodes offered opportunities for gene flow across the EC-MA when lowland ecosystems expanded altitudinally, enhancing dispersal, thus explaining the relatively young divergence times across the Andes as well as the slightly younger median divergence times (estimated by BEAST) for mammals (1.43 Ma) and birds (2.58 Ma) relative to amphibians (3.26 Ma) and in non-avian reptiles (2.99 Ma), as estimated by our phylogenetic gene tree method (Fig. 4).

### Alternative scenarios to Andean vicariance

Our synthesis of recent geological studies (Figs. 2A and 2B) combined with our comparative phylogeographic analyses failed for two reasons to support the traditional model of vicariance mediated by the northern Andes. Geological evidence suggests the EC-MA had already reached significant elevation as early as 38–33 Ma north of ~4°N (see below), and genetic analyses show that divergence was asynchronous within taxonomic classes, except for mammals, as well as among classes, especially the mammals. Here we consider some alternative explanations for variation among our 37 clades in divergence times across the EC-MA. Under a model of pure vicariance caused by a physical barrier, populations on either side start as polyphyletic entities that, in the absence of subsequence migration, will reach reciprocal monophyly at a rate inversely proportional to effective population size (Avise et al. 1983), such that monophyly would be reached 4 times faster in mtDNA than in nDNA (Hudson & Coyne 2002) or even faster (Crawford 2003). Of the 37 tetrapod lineages studied here, 17 (46%; 3 amphibians, 3 non-avian reptiles, 9 birds, and 2 mammals) showed polyphyly with respect to the position of the EC-MA. If this frequent polyphyly is due to incomplete lineage sorting, then this provides further evidence that the hypothesized vicariant split was recent relative to population size, further suggesting that any potential impact of the EC-MA on tetrapods tended to take place more recently than traditionally accepted dates for the end of the uplift of the EC-MA. The asynchrony in timing of divergence makes this explanation less likely, however. Alternatively, polyphyly could be created by multiple crossings of the EC following an initial vicariant separation, as found in, for example, the bird *Mionectes* (Miller et al., 2008) and the frog, *Rheobates* (Muñoz-Ortiz et al., 2015), suggesting that the EC-MA is an ineffective barrier for half of tetrapod lineages studied here. Such a model of recent dispersal across a fully formed Andes could explain the heterogeneity in divergence times among lineages, the overall young ages relative to the old age of the EC-MA as synthesized here, and the young divergence times among homeotherms relative to poikilotherms (Fig. 5).

Our goal here is to synthesize a geological perspective with evolutionary genetic inferences to better understand the biogeographic patterns and processes behind divergence across the northern Andes. As discussed above, the geological evidence suggests a much older and complex EC-MA uplift, yet the genetic data show asynchronous divergence (except in mammals) with median dates that tend to be too young: 3.26 Ma in amphibians, 2.99 Ma in non-avian reptiles, 2.58 Ma in birds, and 1.43 Ma in mammals (Fig. 4). We propose that Pleistocene climate fluctuations facilitating dispersal over the EC-MA. One alternative trans-Andean dispersal scenario, however, could include dispersal through lowland passes. As mentioned in the Introduction, the Magdalena River valley on the Caribbean coast of Colombia contains fossil fishes of Amazonian affinities, suggesting that a pass through the southern EC was open until at least 11 Ma, and fossil catfishes from northern Venezuela suggest a pass existed through the MA as recently as 3–5 Ma (Aguilera et al., 2013). If such connections existed, the geological data is not yet clear where these passages would have been located, but we propose two possible lowland passes through an otherwise tall EC-MA. The western end of the MA could have connected Amazonia and the Caribbean coast to the north ~3 Ma. The junction of the Central Cordillera and southern end of the EC in Colombia, allowed lowland connectivity between Amazonia and the Magdalena ~11 Ma. A potentially younger pass could be represented by the geologically youngest section of the EC (Kroonenberg et al., 1981; van der Wiel, 1991; Ujueta, 1999). Conducting phylogeographic studies across these potential gaps may prove challenging, however, because the environmental conditions today differ dramatically between the two sides, with semi-arid conditions on the western side contrasting sharply with the humid Amazonian moist forest to the East, and very few if any species are found in both.

We report four main observations here that we hope can be accounted for by a single historical explanation: 1) geological evidence revealing old age of uplift for the EC-MA in general dating to around 38–33 Ma, coupled with 2) paleontological evidence from Amazonian lineages of fishes found in the northern Magdalena Valley at 11 Ma and northern Venezuela at 3–5 Ma. 3) asynchrony (aka, high variance) in divergence times (estimated by hABC) within and among groups of tetrapods, ranging from 0.13 Ma in *Boana boans* to 9.05 Ma in *Boa constrictor*, and finally 4) low median divergence times (estimated by BEAST) within each tetrapod group from 3.26 Ma in amphibians to 1.43 Ma in mammals. Because a traditional model of Andean uplift cannot account for the young dates or the variation in divergence times among tetrapod lineages spanning the northern Andes, we propose the following explanation. The significant albeit weak relationship between divergence times and thermal biology (Fig. 5) provides a clue that the interaction between life history and environment may play an important role in structuring variation across the Andes, as has been found in a related comparative phylogeographic study of frogs in Panama (Paz et al. 2015). While the EC-MA was largely formed in the Late Eocene (38–33 Ma), climatic fluctuations continued and were especially strong during the Pleistocene (Flantua et al., 2019). The cooling in the Pliocene and subsequent Pleistocene climate oscillations likely created a ‘flickering connectivity’ (Flantua et al., 2019) between lowland populations east and west of the EC-MA, and the connectivity would have been greater in homeotherms relative to poikilotherms since the former can withstand lower temperatures and thus disperse more easily across an elevational gradient. Thus, environmental variation, not geology, recently separated lineages on either side of the EC-MA.

Geology and evolutionary genetics offer complementary views of the history of our planet. Any historical hypothesis generated from genetic data should be evaluated with geological information, and vice versa. The present analysis is unique in explicitly synthesizing geological and genetic perspectives, including one of the first comparative studies involving organisms with widely varied physiological and ecological traits, to evaluate the role of the northern end of the World’s longest mountain chain, the Andes of South America, in promoting lowland diversification. We conclude that mountains did not limit dispersal, climate did, and we look forward to further testing of our hypothesis using a greater diversity of lineages and genetic markers.

## Acknowledgments

For collecting permits, we thank the *Ministerio de Ambiente y Desarrollo Sostenible* and the *Autoridad Nacional de Licencias Ambientales* (ANLA) in Colombia (*Permiso de estudio con fines de investigación científica en diversidad biológica No. 27 del 22 de junio de 2012*, *permiso de acceso a los recursos genéticos resolución No. 0377 del 11 marzo de 2014* to Andrew J. Crawford (A.J.C.) and *permiso marco resolución No. 1177* to the Universidad de los Andes). We thank Santiago Castroviejo-Fisher for providing sequences of *Dendropsophus microcephalus, Boana pugnax, Elachistocleis*, and *Scarthyla vigilans*. We are very grateful to Michael J. Hickerson for providing feedback and recommendations for running MTML-msBayes. This project was funded by the *Facultad de Ciencias* of the Universidad de los Andes under the call *Proyectos cortos o generación de un producto adicional*, and DIDI (*Dirección de Investigación, Desarrollo e Innovación*) of Universidad del Norte. New DNA sequence data reported here for frogs were obtained with the generous support of a grant in Basic Sciences No. 360-2013 from *Colciencias* to AJC.

## Biosketches

**Erika Rodríguez-Muñoz** is interested in biogeography and evolutionary ecology of plants and animals in the Neotropics.

**Camilo Montes** is a structural geologist interested in learning about paleogeography and how different tectonic configurations through time may have impacted climate, global circulation, and life.

**Andrew J. Crawford** (http://www.dna.ac/) is an evolutionary geneticist interested in the history and mechanisms of speciation and adaptation in frogs and other tetrapods in the American Tropics. Find him on Twitter via @CrawfordAJ

## Author contributions

Authors jointly conceived the ideas. C.M. synthesized the geological information. A.J.C. supervised collection of the genetic data for frogs but did not actually pick up a pipettor. E.R.M. did the real work of organizing the previously published DNA data and performing all evolutionary genetic analyses. All authors participated in the writing.

